# Exploring priority and year effects on plant diversity, productivity and vertical root distribution: first insights from a grassland field experiment

**DOI:** 10.1101/2023.11.14.566982

**Authors:** Inés M. Alonso-Crespo, Vicky M. Temperton, Andreas Fichtner, Thomas Niemeyer, Michael Schloter, Benjamin M. Delory

## Abstract

1. The order of arrival of plant species during community assembly can affect how species interact with each other. These so-called priority effects can have strong implications for the structure and functioning of plant communities. However, the extent to which the strength, direction, and persistence of priority effects are modulated by weather conditions during plant establishment (‘year effects’) is not well known.
2. Here we present the first results from a long-term field experiment (POEM: PriOrity Effects Mechanisms) initiated in 2020 in Northern Germany to test how plant functional group (PFG) order of arrival and the year of initiation of an experiment interactively affect the structure and functioning of nutrient-poor dry acidic grasslands, both above and belowground. To do this, we established the same experiment, manipulating the order of arrival of forbs, grasses and legumes on the same site, but in different years.
3. We found that time since establishment was a stronger driver of plant community composition than PFG order of arrival and year of initiation. These three factors interactively affected plant species diversity, with the effect of PFG order of arrival on plant species richness depending on time since establishment. Year of initiation, not PFG order of arrival, was the strongest driver of aboveground community productivity. Although we did not find any effect of PFG order of arrival on root productivity, it had a strong impact on the vertical distribution of roots. Communities where grasses were sown first rooted more shallowly than communities in which forbs or legumes were sown first.
4. *Synthesis*: Our results demonstrate that plant order of arrival and year effects jointly affect plant diversity and species composition, with time since establishment also playing an important role. While year effects were more important than plant order of arrival in modulating aboveground biomass production in our nutrient-poor grassland, we showed that plant order of arrival can strongly affect the vertical distribution of roots, with communities in which forbs or legumes were sown first rooting deeper than grasses-first communities. These results suggest that a deeper understanding of priority and year effects is needed to better predict restoration outcomes.

## Introduction

Grasslands are a dominant land use type all across the globe. Their conservation and restoration are critical to combat both the biodiversity and climate crises. In the UN Decade on Ecosystem Restoration (2021-2030), the pivotal role that species-rich grasslands could potentially play in both ensuring resilience in the face of extreme weather events, storing belowground carbon (Dass et al., 2018) as well as swiftly restoring biodiversity (Staude et al., 2023; Temperton et al., 2019) is gaining traction. Against this backdrop, there is increasing interest in implementing effective multifunctional grasslands into existing landscapes (Temperton et al., 2019; Wilsey, 2020). Until recently, however, species-rich grassland restoration was mainly focused on creating as biodiverse grasslands as possible, with aspects of ecosystem functioning (such as productivity, nutrient cycling or carbon storage) having only attracted concerted attention within more academic biodiversity-ecosystem functioning (BEF) experiments (Lambers et al., 2004; Weisser et al., 2017). Thus, we still lack a comprehensive understanding of how to restore grasslands to promote habitat-specific plant diversity and enhance ecosystem functioning at the same time.

Grassland experiments that manipulate community assembly such that different plant functional groups (hereafter called PFG) arrive earlier than others have shown that the legacy of which functional group establish first at a site after disturbance can alter not only plant diversity but also ecosystem productivity (Delory et al., 2019; Körner et al., 2008; von Gillhaussen et al., 2014; Weidlich et al., 2017). Such so-called priority effect approaches, whereby arrival order or initial relative abundance of species can significantly influence further community assembly, community structure and composition and/or ecosystem functioning (Drake, 1991; Fukami, 2015) are well known within the field of ecological restoration (Funk et al., 2008). Many priority effect studies have focused on reducing the performance of unwanted invasive species by sowing target species first to keep the invader out (Hess et al., 2019; Martin & Wilsey, 2012; Yannelli et al., 2020). However, the key limitation for grassland restoration is not invasive species, but a need for restoring diversity and aiming for higher productivity, resilience to drought or sequestering carbon in soils (Lyons et al., 2023). Sowing certain species or PFG before others may induce desired trajectories that create the kind of multifunctional outcomes we are currently looking for (Weidlich et al., 2021).

Studies that have followed the PFG approach cluster species based on their traits or degree of phylogenetic relatedness, and manipulate their order of arrival in grassland systems under controlled and field conditions (see Weidlich et al., (2021) for an overview). These studies found clear effects of PFG order of arrival on productivity, both aboveground (Körner et al., 2008; von Gillhaussen et al., 2014; Weidlich et al., 2017) and belowground (Körner et al., 2008; Weidlich et al., 2018), with communities where legumes were sown before the rest often having higher aboveground productivity, but lower root biomass in the topsoil. In a controlled experiment, we showed that manipulating PFG order of arrival can also affect the vertical distribution of roots in the soil, with communities where grasses were sown first rooting more shallowly than communities where either forbs or legumes were sown first (Alonso-Crespo et al., 2023). It is not known whether these results are also valid under field conditions, which argues in favor of longer-term experiments to monitor root development and distribution, using minirhizotron tubes for example.

We expect stronger priority effects in ecosystems with higher availability of resources (Chase, 2003), since in more nutrient-rich sites competitive species may be better able to preempt resources and thus affect later arriving species through asymmetric competition, creating stronger priority effects. Conversely, if such priority effects include facilitative interactions, this may lead to more ecologically even communities that can create both plant diversity as well as perhaps resilience, better carbon storage and increased hay productivity (as a motivation to farmers to maintain such grasslands). Previous PFG priority studies have generally focused on more mesotrophic grassland settings (Weidlich et al., 2017), whereas the current study examines whether we can find above and belowground priority effects in a low nutrient dry acidic grassland.

Despite the importance of time and persistence of priority effects for trajectories and alternative states of plant communities, little attention has so far been paid to either long term or year effects (e.g. the environmental conditions in which the initial community assembly takes place) possibly since it requires a huge effort to set up the same experiment repeatedly over time (Stuble et al., 2017.b). Evidence for year effects has been hard to extract from studies due to correlational considerations and confounding factors (Groves et al., 2020; MacDougall et al., 2008), but Stuble et al. (2017.a) studied the effect of year of initiation and site on restoration outcomes and found both strong site and year effects on community composition. In another study, Stuble et al. (2017.b) manipulated the timing of arrival of native and exotic grasses in Californian grasslands across different sites and years, and found that the strength of priority effects was strongly dependent on the location of the experimental site and the year in which an experiment was initiated. A study by Groves & Brudvig (2019) where year of initiation and precipitation were manipulated underlined that year effects can occur even if precipitation is held constant, suggesting an important role for environmental variables other than precipitation in driving year effects during restoration. Werner et al. (2020) went further by including two different sowing intervals (2 weeks and one year) within priority effect treatments across sites and initiated in different years. They found that the year effect was by far the largest driver of outcomes, followed by priority (sowing interval) and site treatments. Interannual variation in environmental conditions during plant establishment has also been shown to be an important driver of the taxonomic and functional composition and diversity of plant communities, with consequences for ecosystem functioning (Werner et al., 2020; Atkinson et al., 2023; Catano et al., 2023). To better predict restoration outcomes, it is necessary to delve deeper into the ecological mechanisms underlying year effects and their context-dependence.

Although it has been shown that PFG order of arrival can affect community structure, above and belowground productivity, and root distribution in the soil (Alonso-Crespo et al., 2023; Körner et al., 2008; von Gillhaussen et al., 2014; Weidlich et al., 2017, 2018), little is known about how persistent these priority effects are over time and how this is mediated by the environmental conditions during establishment. Additionally, belowground, it remains uncertain whether priority effects on root productivity are a consequence of changes in the total root productivity or changes in the vertical root distribution.

In this study, we present the first results of a long-term field experiment (POEM, <u>P</u>ri<u>O</u>rity <u>E</u>ffect <u>M</u>echanisms) initiated in 2020 and designed to test how priority and year effects modulate the structure and functioning of dry acidic grassland plant communities over time, both aboveground and belowground. To do this, we set up independent sub-experiments at a site in Northern Germany, where we tested the same PFG order of arrival scenarios, but in different years. Here, we used species-specific shoot biomass data collected between 2020 and 2023, as well as root images taken at different depths using minirhizotrons between 2021 and 2023, to address the following hypotheses:

1. Manipulating PFG order of arrival affects species composition and plant diversity, with plant communities following different trajectories depending on the year of initiation of an experiment.
2. PFG order of arrival and year of initiation interactively affect the aboveground productivity of plant communities. Following previous work (Körner et al., 2008; Weidlich et al., 2017), we expect plant communities in which legumes were sown first to be the most productive, but not necessarily for each year of initiation.
3. Root productivity in the first 50 cm of soil depends on PFG order of arrival, with plant communities in which legumes were sown first being the least productive belowground (Körner et al., 2008; Weidlich et al., 2018).
4. PFG order of arrival affects the vertical distribution of roots at the community level. Following previous work (Alonso-Crespo et al., 2023), we expect communities in which forbs or legumes were sown first to root deeper than communities in which grasses were sown first.

## Material and Methods

### Study site

The POEM experiment is located on a former arable land owned by a local conservation organisation (Verein Naturschutzpark, VNP) in a fenced area in Niederhaverbeck, Germany (latitude: 53.144272, longitude: 9.912668; altitude: 105 m a.s.l.; mean annual air temperature: 10.2°C; minimum air temperature: −14.9°C; maximum air temperature: 38.7°C; mean annual precipitation: 684 mm, 2020-2023). The experiment was set up on a soil that is appropriate for establishing dry acidic grassland communities (sand fraction: 93%; clay and silt fraction: 4%; pH (CaCl_2_) 4.9; organic matter content: 2.3%; total N: 0.07%; total C: 0.98%; C/N: 12.1). Our experimental site had been used during the last 200 years as a cultivated arable field. In the years preceding the experiment, the following species were grown in our experimental area: *Trifolium repens*, *Trifolium pratense*, *Trifolium incarnatum*, *Lolium perenne*, Festulolium (*Festuca* sp. × *Lolium* sp.) and *Secale cereale*.

### Species pool and classification into plant functional groups

A total of fourteen plant species were used in this field experiment: four N_2_-fixing legumes (*Lathyrus pratensis* L., *Lotus corniculatu*s L., *Trifolium arvense* L., and *Trifolium campestre* Schreb.), four grasses (*Agrostis capillaris* L., *Anthoxanthum odoratum* L., *Bromus hordeaceus* L., and *Festuca ovina* agg.), and six forbs (*Dianthus deltoides* L., *Jasione montana* L., *Pilosella officinarum* L., *Pimpinella saxifraga* L., *Potentilla argentea* L., and *Silene vulgaris* (Moench) Garcke). These species were chosen based on typical plant functional group ratios found in dry acidic grasslands (i.e. more forbs than grasses and legumes), as well as their availability from a regional/local wild seed company. The seeds of all species were obtained from Rieger-Hofmann GmbH (Blaufelden, Germany).

### Experimental design

The POEM field experiment was set up using a full factorial design to test for the influence of (1) plant functional group (PFG) order of arrival and (2) the year of initiation of an experiment on the aboveground and belowground structure and functioning of dry grassland plant communities.

Year of initiation effects are tested by setting up the same experiment at the same site, but in a separate block, in four different years. Here, we use the results obtained for the first two sub-experiments set up in 2020 (referred to as POEM2020) and 2021 (referred to as POEM2021), respectively. Two additional sub-experiments will be set up in the coming years.

In each POEM sub-experiment, we manipulated the order of arrival of forbs, grasses and legumes. We tested five arrival scenarios: (1) simultaneous sowing of forbs, grasses and legumes at the first sowing event (synchronous, S), (2) forbs sown six weeks before grasses and legumes (F), (3) grasses sown six weeks before forbs and legumes (G), (4) legumes sown six weeks before forbs and grasses (L), and (5) no sowing of additional species (free succession, B) (Figure 1). Each arrival scenario was replicated 5 times in each sub-experiment. Thus, each POEM sub-experiment consists of 25 mixture plots of 9 m^2^ (3 m×3 m). Next to these mixture plots, we also set up 14 monoculture plots of 4 m^2^ (2 m×2 m) in which each species from our species pool grew on its own and here the plots were regularly weeded. New monoculture plots were set up for every year of initiation. Within each sub-experiment, experimental treatments and species identity were randomly assigned spatially to mixture and monoculture plots, respectively. Both PFG order of arrival (5 levels: S, F, G, L, B) and the year of initiation of the experiment (2 levels: POEM2020 and POEM2021) are considered fixed factors in the experiment.

**Figure 1.**
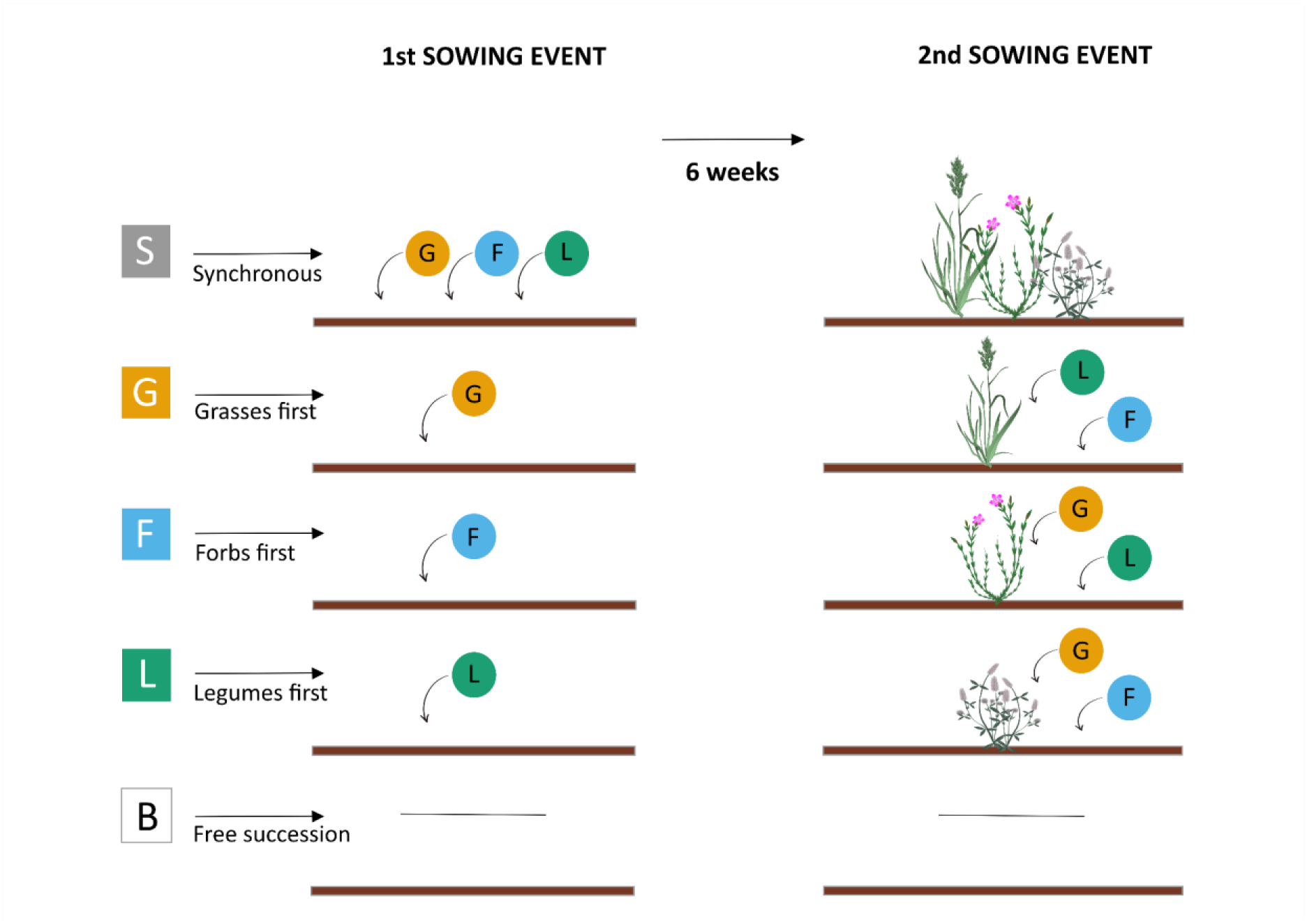
PFG order of arrival scenarios tested in the first two POEM sub-experiments. We tested five PFG order of arrival scenarios: (1) simultaneous sowing of forbs, grasses and legumes at the first sowing event (synchronous, S), (2) forbs sown six weeks before grasses and legumes (F), (3) grasses sown six weeks before forbs and legumes (G), (4) legumes sown six weeks before forbs and grasses (L), and (5) no sowing of additional species (free succession, B).

Our experimental site has been equipped with a weather station that continuously monitors air temperature (°C), air relative humidity (%), precipitation (mm), precipitation intensity (mm/h), wind speed (m/s), wind direction (°), atmospheric pressure (hPa), global radiation (W/m^2^), photosynthetically active radiation (µmol m^−2^ s^−1^), soil volumetric water content at 10, 20, 30, and 40 cm (%), and soil temperature at 5 cm (°C) (Figure S1.A).

### Experimental setup

Before starting each sub-experiment, the experimental area was harrowed, plots were marked and weeded, and large stones were removed by hand. The first sowing event took place on April 27, 2020 (for POEM2020) and April 13, 2021 (for POEM2021). The second sowing event took place six weeks later on June 8, 2020 (for POEM2020) and May 25, 2021 (for POEM2021) (Figure S1.B). Before starting each sub-experiment, the germination rate of each species was measured under controlled conditions. Mixture and monoculture plots were sown with 1000 viable seeds per m^2^. Seed mixtures were prepared so that (1) all PFG had the same relative abundance and (2) all species within each PFG had the same relative abundance. A detailed description of the composition of the seed mixtures used for each sub-experiment is provided in Tables S1 and S2. In the first sowing, the seeds were mixed with two cups of sand and spread evenly over the surface of the plots. A similar approach was used in the second sowing, but the seeds were sown from above, as we did not mow the plots before the second sowing and we were careful not to disturb the plants growing in the plots. After starting each sub-experiment, we stopped weeding unsown species (i.e., species invading the plots or originating from the soil seedbank) because the longer term goal is to simulate ecological restoration where only one (or two) sowing events usually are used and then the site is allowed to undergo natural assembly processes. Monoculture plots, however, were regularly weeded. The surroundings of the experiment and the paths between the plots were sown with the non-clonal grass *Festuca rubra* spp. *commutata* Gaudin. All plots were mown once a year in August.

### Installation of minirhizotron tubes

In order to be able to non-destructively monitor root development and distribution over time, we installed acrylic minirhizotron tubes in synchronous, forbs-first, grasses-first, and legumes-first plots of POEM2021. We did this in October 2020, i.e. six months before starting the second POEM sub-experiment. In each plot, two one-metre long minirhizotron tubes (Vienna Scientific Instruments Gmbh; outer diameter: 60 mm; wall thickness: 3 mm) were installed at a 45° angle using a 58-mm diameter soil corer (Vienna Scientific Instruments Gmbh). All minirhizotron tubes were closed on both sides and were installed as shown in Figure S2, with the upper 21 cm of the tubes sticking out of the soil. The aboveground portion of the observation tubes was closed with a water- and light-tight plastic cap and covered with pipe insulation foam to exclude light and reduce thermal fluctuations. Within a plot, minirhizotron tubes were installed in the same direction (east-west or west-east), which was randomly selected for each plot. Between October 2020 and April 2021, the experimental area of POEM2021 was covered with a water-permeable ground sheet in order to avoid plant growth. A step-by-step description of the installation of minirhizotron tubes in POEM2021 is available in the video provided as supplementary material.

### Aboveground data collection

The structure of plant communities was assessed once a year by recording the total shoot dry weight of each individual species (sown and unsown species) located inside two randomly positioned 0.1 m^2^ quadrats (20 cm × 50 cm). Four quadrats per plot were harvested at the end of the first growing season of POEM2020, but data showed that good estimates of community productivity can be achieved with only two quadrats. These harvests were organised each year during the peak biomass production in June/July. At harvest, plants were cut 3 cm above the soil surface and sorted at the species level directly in the field. Shoot samples were dried in an oven at 60°C for at least 48 h and weighed on an analytical scale. For each plot, total aboveground productivity was calculated as the sum of the contributions of each individual species growing in the communities.

In order to get better estimates of the realised plant species richness inside the plots, the presence and absence of sown and unsown species was also monitored once a year, typically before collecting shoot biomass samples. These data were collected in all sub-experiments as of June 2021.

### Root image acquisition

High-resolution images of roots growing along the transparent minirhizotron tubes were acquired regularly using the VSI MS-190 manual camera (Vienna Scientific Instruments Gmbh). Root images (2340×2400 pixels; resolution: 148 pixels/mm) were taken at 18 different depths, equally spaced from 1.4 cm to 49.5 cm deep (Figure S2). When taking root images, the camera was always pointing upwards. Weather permitting, image acquisition was conducted twice a month from April 2021 to September 2021, once a month from October 2021 to March 2022, twice a month from April 2022 to September 2022, and once a month thereafter. Images were collected at 33 time points between April 2021 and June 2023 (3 growing seasons), which represents a total of 23,720 root images.

### Root image analysis

Root images were analysed using an approach similar to (Alonso-Crespo et al., 2023). First, we used RootPainter (Version: 0.2.27) to train a convolutional neural network to detect roots in our images (Smith et al., 2022). To do this, we first created two independent datasets of 1,440 images. Each dataset consisted of all the images taken at two different time points, one year apart. We did this to ensure that each dataset included a wide variety of images. Each dataset consisted of images from different time points. Then, we used RootPainter to randomly select two subregions (800×800 pixels) from each image of each dataset, thus creating two training datasets of 2,880 images each. Each training dataset was then assigned to a different user with previous experience with image annotation and model training with RootPainter. Each user annotated at least 500 images of their training dataset (corrective annotations) and trained their model until predictions successfully identified most of the roots in training images. Both users used the same set of rules for annotating images. In particular, users aimed to train a model able to detect the centerline of living roots, thus avoiding dead roots, root edges, soil background, water droplets and scratches at the surface of the minirhizotron tubes. This strategy proved useful in segmenting roots growing next to each other separately. Following this initial training procedure, the two users combined their training images and annotations and began training a third model, using the best of their two models as a starting point. This third model performed better than the models obtained by each user alone and was used to segment all root images in our dataset. Model training and image segmentation were done on a GPU node (NVIDIA Ampere A100 GPU with 40 GB memory) of a computer cluster available at the Leuphana University Lüneburg (Germany).

Segmented images were analysed with RhizoVision Explorer 2.0.3 (Seethepalli et al., 2021) to estimate the total root length and the projected root surface area in each image. These parameters were then used to estimate the planar root length density (*pRLD*, total root length divided by image area, cm cm^−2^) and the planar root surface density (*pRSD*, projected root surface area divided by image area, cm^2^ cm^−2^). Both *pRLD* and *pRSD* were highly correlated (r=0.96, P < 0.0001). Because our image segmentation model was trained to detect the centerline of the roots, we probably underestimated *pRSD* and decided to focus on *pRLD* in this paper (but see supplementary information for results using *pRSD*).

## Data analysis

Differences in plant community structure and community assembly trajectories were visualised using non-metric multidimensional scaling (NMDS). NMDS was performed with the metaMDS function of the R package vegan (Oksanen et al., 2022). We performed this analysis twice, on two different datasets: (1) species-specific plant biomass data (continuous data) measured for the three first growing seasons of POEM2020 and POEM2021, and (2) species presence/absence (binary data) measured as of June 2021. We used the Bray-Curtis dissimilarity index for plant biomass data and the Jaccard dissimilarity index for presence/absence data. Before NMDS, square root transformation and Wisconsin double standardisation were applied to plant biomass data. In order to test for differences in composition between treatments, we conducted two permutational multivariate analyses of variance (PERMANOVA) using PFG order of arrival (categorical variable with 5 levels: S, F, G, L, B), year of initiation (categorical variable with 2 levels: POEM2020 and POEM2021), sampling year (categorical variable with 3 levels: year 1 to 3) and their interactions as fixed factors (Anderson, 2001). F-statistics and P-values were computed based on 1000 permutations. All PERMANOVA models were fitted using the adonis2 function of the R package vegan (Oksanen et al., 2022). The same distance matrix was used for the NMDS and the PERMANOVA. Although distance-based multivariate analyses are known to confound location and dispersion effects (Warton, Wright & Wang, 2012), PERMANOVA has been shown to be largely insensitive to heterogeneity in multivariate dispersion between groups in the case of balanced designs (Anderson & Walsh, 2013), which is the case in this study.

The effects of PFG order of arrival, year of initiation and sampling year on plant diversity were analysed at the alpha level using Hill numbers (Chao et al., 2014). For each combination of experiment, sampling year and PFG order of arrival, we constructed diversity profile plots to visualise how the effective number of plant species (*D*) changes with diversity order (*q*). Diversity order is a parameter used to adjust the sensitivity of *D* to the relative abundance of species in a community. We calculated effective taxonomic diversity (*D*) of order *q* for *S* species using Equations 1a (*q* ≥ 0, *q* ≠ 1) and 1b (*q* = 1), where *p_ij_* is the relative abundance of species *i* in plot *j*. This was done using the hillR R package (Li, 2018). When *q*=0, all species are weighted equally, which is equivalent to species richness. Using larger values of *q* increases the weight of abundant species relative to rare ones. *^1^D* (i.e., effective diversity of order 1) is equivalent to the exponential of Shannon entropy, while *^2^D* (i.e., effective diversity of order 2) is equivalent to the inverse Simpson concentration. In terms of interpretation, *^q^D = x* means that the diversity of order *q* of this assemblage is equivalent to an idealised assemblage consisting of *x* equally abundant species. The effects of PFG order of arrival, year of initiation, sampling year and their interactions on plant diversity at *q*=0, *q*=1 and *q*=2 were analysed using three separate generalised linear mixed-effect models. Each model was fitted with the lme4 R package (Bates et al., 2015) using a Gamma distribution and a log-link function. Because data were collected in the same plots for three years, plot ID was used as a random effect in the model (random intercept). Post-hoc tests were carried out using the emmeans R package (Lenth, 2023).

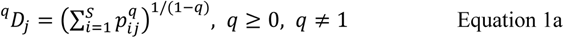

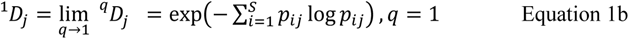

The effects of PFG order of arrival, year of initiation, sampling year and their interactions on total aboveground productivity (g/m^2^) were analysed using a generalised linear-mixed-effect model. Plot ID was used as a random effect (random intercept). The model was fitted with the lme4 R package (Bates et al., 2015) using a Gamma distribution and a log-link function. Post-hoc tests were carried out using the emmeans R package (Lenth, 2023).

As a proxy for root productivity in a plot at a given time point, we calculated the average planar root length density (*pRLD*) across 36 minirhizotron images (2 tubes/plot × 18 images/tube). We then modelled the temporal evolution of *pRLD* for each PFG order of arrival using a generalised additive model (GAM). The model included three components: (1) a fixed effect for PFG order of arrival, (2) a smooth function of time since the start of the experiment (thin plate regression spline), conditioned by PFG order of arrival (the number of basic functions, *k*, was set to the number of time points in the dataset), and (3) a random effect smooth for plot ID. This model was fitted with the mgcv R package (Wood, 2017) using a tweedie family distribution with a log-link function. We used the same approach to model the temporal evolution of *pRSD* for each PFG order of arrival.

We assessed how PFG order of arrival affected the vertical distribution of roots in two complementary ways: (1) by modelling the temporal evolution of the average rooting depth of plant communities, and (2) by modelling the evolution of *pRLD* as a function of time and soil depth. The same approach has been used for *pRSD* data, so we focus solely on *pRLD* in the following paragraphs.

At each time point, the mean rooting depth (*MRD*) in plot *j* was calculated using equation 2, where *d_ij_* is the soil depth at location *i* in plot *j*, and *pRLD_ij_* is the average *pRLD* measured at location *i* in plot *j*. The temporal evolution of *MRD* was then modelled using a generalised additive model consisting of three components: (1) a fixed effect for PFG order of arrival, (2) a smooth function of time since the start of the experiment (thin plate regression spline), conditioned by PFG order of arrival (the number of basis functions, *k*, was set to the number of time points in the dataset), and (3) a random effect smooth for plot ID. This model was fitted with the mgcv R package (Wood, 2017) using a tweedie family distribution with a log-link function.

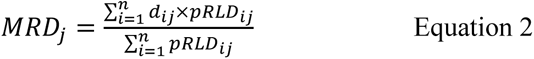

Changes in *pRLD* as a function of time and soil depth were modelled using a generalised additive model. The model included three components: (1) a fixed effect for PFG order of arrival, (2) a tensor product smooth function of time since the start of the experiment and soil depth (cubic regression spline), conditioned by PFG order of arrival, and (3) a random effect smooth for plot ID. This GAM was fitted with the mgcv R package (Wood, 2017) using a tweedie family distribution with a log-link function.

Data analysis was performed in R version 4.2.3 (R Core Team, 2023). In addition to the R packages mentioned above, the following R packages were used for data exploration, visualisation and analysis: R packages included in tidyverse (Wickham et al., 2019), car (Fox & Weisberg, 2019), Hmisc (Harrell, 2023), ggpubr (Kassambara, 2023), gtools (Bolker, Warnes & Lumley, 2022), ggConvexHull (Martin, 2017) and viridis (Garnier et al., 2021).

## Results

### Time since establishment drives plant community composition more strongly than PFG order of arrival and year of initiation

Overall, time since establishment (partial R^2^: 37-39%) had a much stronger effect on the composition of plant communities than PFG order of arrival (partial R^2^: 8-10%) and the year of initiation of an experiment (partial R^2^: 6-8%). This result was consistent for both biomass (Figure 2A, Table S3) and presence/absence data (Figure 2B, Table S4). The effect of PFG order of arrival on species composition was similar across combinations of year of initiation and sampling year (Figure 2A, P = 0.082; Figure 2B, P=0.966). Except for free succession plots, plant communities followed similar, but not identical, trajectories across PFG order of arrival scenarios within each sub-experiment, confirming that PFG arrival order plays a role, albeit a modest one, in modulating species composition (see Tables S3-S4 and next section on plant diversity). Plant communities in both experiments remained different over the 3-year study period, which highlights the important role of year of initiation for community assembly in our POEM experiment (Figure 2, Tables S3-S4).

**Figure 2.**
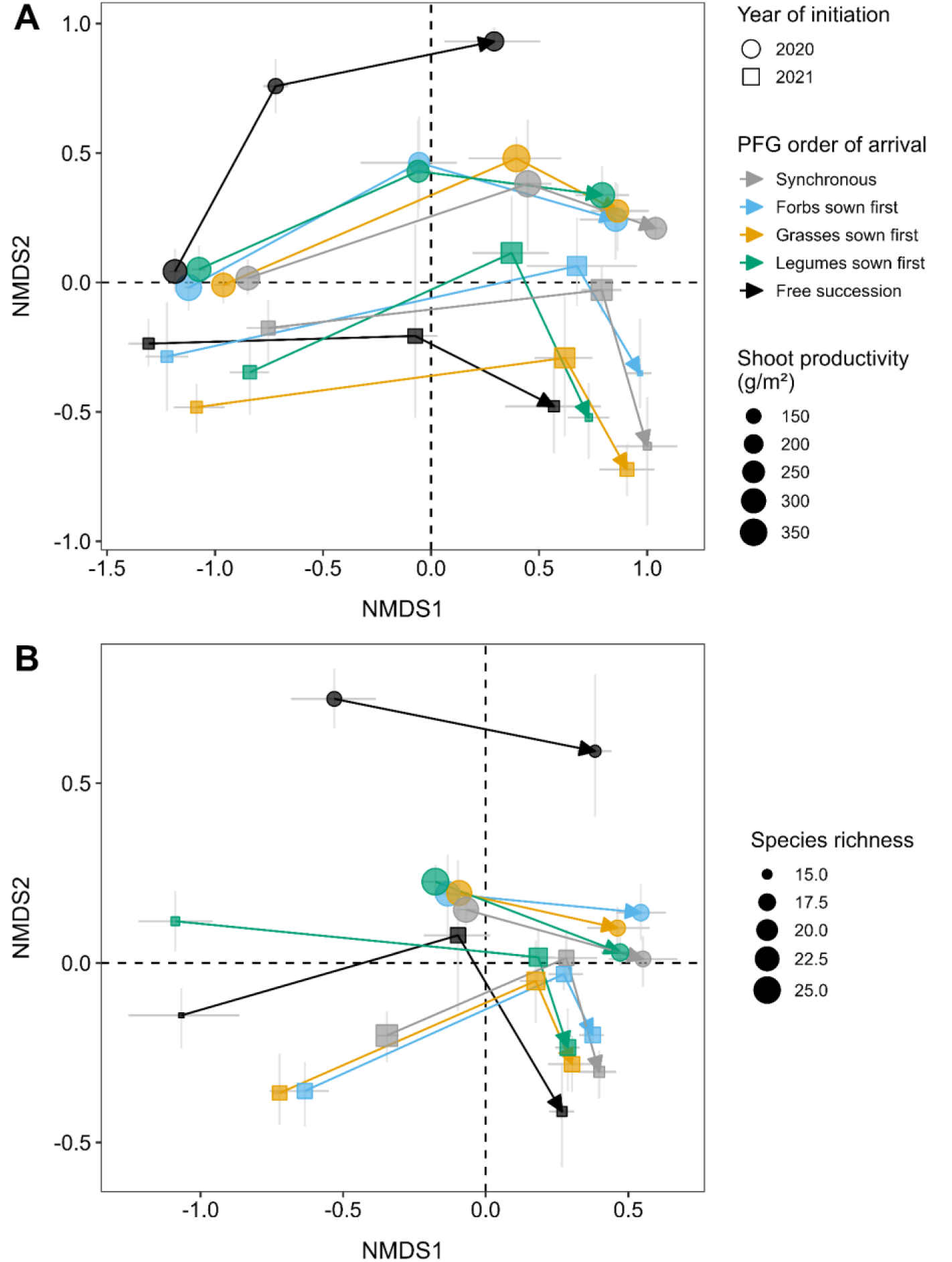
Time since establishment drives plant community composition more strongly than PFG order of arrival and year of initiation. Panels A and B are non-metric multidimensional scaling (NMDS) plots showing dissimilarities in plant community composition observed after one, two and three growing seasons between plots of two POEM sub-experiments (POEM2020 and POEM2021) in which PFG order of arrival was manipulated. Species-specific biomass data were used in panel A (stress: 0.166). Species presence/absence data were used in panel B (stress: 0.148; no data available for the first growing season of POEM2020). For each combination of year of initiation, sampling year and PFG order of arrival, the centroid position and associated 95% confidence intervals computed using non-parametric bootstrap (gray segments) are shown (n=5).

During the first growing season of both sub-experiments, plant communities were strongly dominated by unsown species, which represented on average 97% and 93% of the total amount of biomass collected at harvest in POEM2020 and POEM2021, respectively. It was not until the second growing season that the sown species started to take over in the treatment plots, particularly in POEM2021 (Figure S3). In the second growing season, sown species represented on average 77% and 93% of the harvested biomass in the first and second sub-experiments, respectively (Figure S3). The relative proportion of sown species measured at the end of the third growing season reached very similar values (72% for POEM2020, 93% for POEM2021).

In both sub-experiments, forbs were most productive in plots where this PFG was given a head start, while grasses (especially *Bromus hordeaceus*) were usually more abundant in synchronous and grasses-first plots, particularly at the end of the second growing season (Figure S4). Legumes, however, showed a more complicated pattern. Although *Trifolium arvense* established quite well in synchronous and legumes-first plots of POEM2021, legumes were not necessarily the most productive in plots where they were sown first (Figure S4).

### Time since establishment, PFG order of arrival and year of initiation interactively modulate plant diversity

PFG order of arrival had distinct effects on plant diversity in each sub-experiment, particularly in the two first growing seasons (Figure 3, see Tables S5-7 for statistics). Plant species richness (*q*=0) was weakly affected by the year of initiation of an experiment, but was mainly dependent on PFG order of arrival and time since establishment (Figure 3, Table S5). At the end of the first growing season, we harvested on average seven species more in synchronous plots than in free succession plots in both sub-experiments. In synchronous communities of POEM2021, we also harvested on average four to six species more than in plots where one PFG was sown before the other two. When more weights are given to abundant species (i.e., at higher values of *q*), plant diversity decreased and did not differ between PFG arrival scenarios, except for forbs-first plots which were less diverse than synchronous plots at the end of the first growing season of POEM2021. When more weights are given to very dominant species (*q*=2), our plant communities were equivalent to an idealised assemblage with 3 (POEM2020) and 2 (POEM2021) equally abundant species, which is due to the strong dominance of the unsown species *Spergula arvensis* (58%), *Erodium cicutarium* (13%), *Anthemis arvensis* (12%) and *Chenopodium album* (6%) after one growing season (Figure 3).

**Figure 3.**
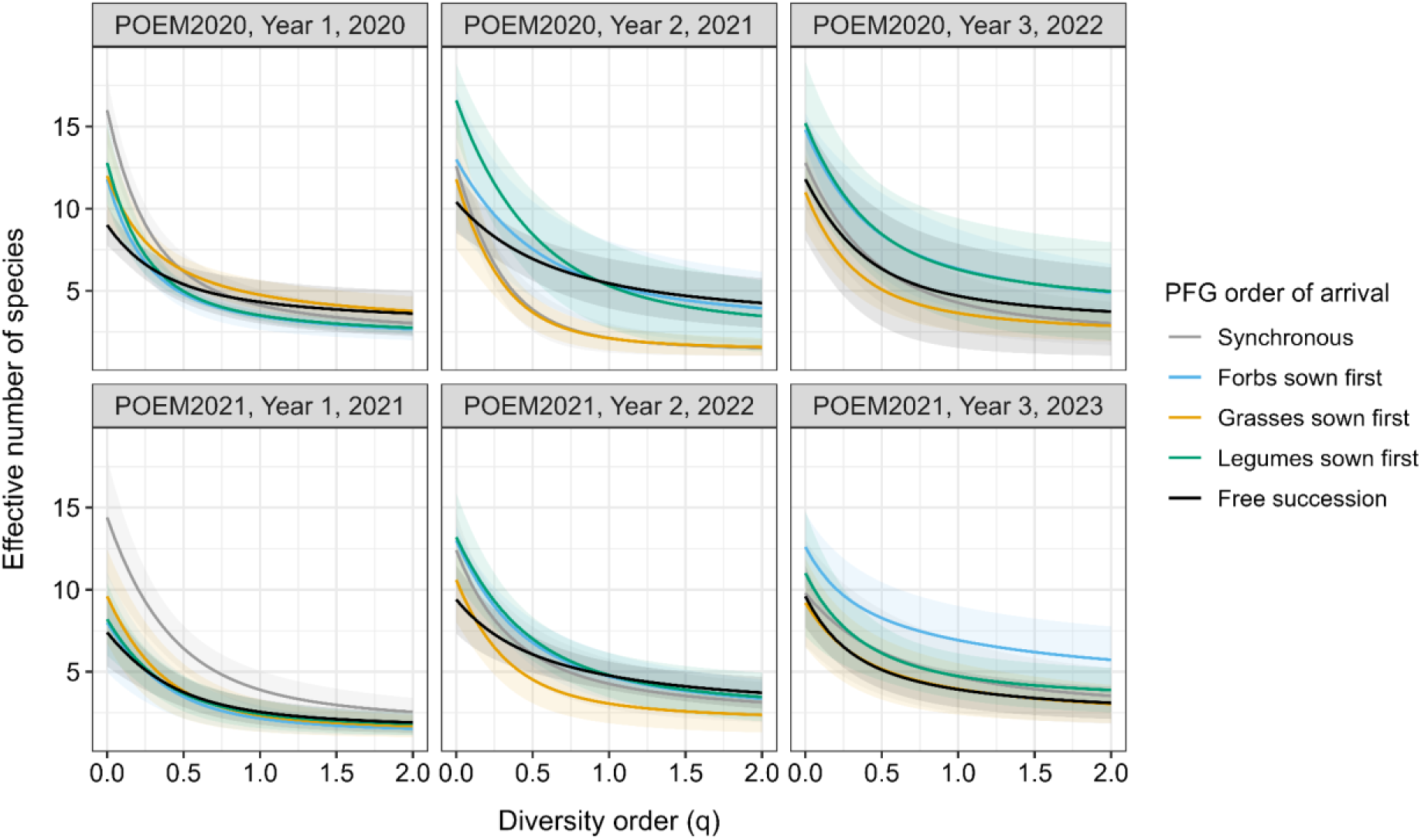
Time since establishment, PFG order of arrival and year of initiation interactively modulate plant diversity. These diversity profile plots show how the effective number of species (*D*) changes as a function of diversity order (*q*) for each combination of year of initiation, sampling year and PFG order of arrival. Lines and shaded areas show the average value and 95% confidence intervals (Gaussian) across five replicates, respectively. *D* values at *q*=0, *q*=1 and *q*=2 are equivalent to species richness, the exponential of Shannon entropy, and the inverse Simpson concentration, respectively. Increasing the value of *q* increases the weight of abundant species relative to rare ones. In terms of interpretation, if *D*=10 at *q*=0, this means that 10 species were present in the community (species richness). If *D*=3 at *q*=2, this means that the species diversity of order 2 of the community is equivalent to an idealised assemblage consisting of 3 equally abundant species (which would suggest that the assemblage contains 3 very dominant species). An assemblage whose species are all equally abundant (i.e., perfectly even assemblage) would be represented by a horizontal diversity profile (i.e., *D* equals to species richness for all values of *q*).

After two growing seasons, we harvested on average four to seven species more in legumes-first plots than in free succession plots in both sub-experiments (Figure 3). Plant species richness, however, did not differ between plots where legumes were sown first and other PFG arrival scenarios, except for grasses-first plots in POEM2020 which had on average five species less than legumes-first plots. With greater weights for abundant (*q*=1) and dominant (*q*=2) species, plant diversity decreased more strongly in synchronous and grasses-first plots in POEM2020, which highlights the overall negative effect that an early arrival of grasses had on plant diversity and community evenness in our experiment. This is mainly due to the strong dominance of the sown grass species *Bromus hordeaceus* (70-81%) in synchronous and grasses-first plots after two growing seasons. We did not find this pattern in our second sub-experiment, although grasses-first plots tended to be on average less diverse than the others (Figure 3).

Interestingly, towards the end of the third growing season, plots in which forbs were sown first became the most diverse (on average, four additional species collected at harvest) and had two times more common (*q*=1) and dominant species (*q*=2) than grasses-first plots (Figure 3). This result was observed in both sub-experiments. In POEM2020, we also found that legumes-first plots were more diverse than grasses-first plots. In POEM2021, plots in which forb species were sown first also contained two times more common and dominant species than free succession plots (Figure 3).

These results indicate that PFG order of arrival modulates plant diversity, but that this effect evolves over time and depends on the year of initiation of an experiment.

### Year of initiation, not PFG order of arrival, is the main driver of aboveground productivity

Across sub-experiments and sampling years, we never found any difference in standing shoot biomass production between plots in which PFG order of arrival was explicitly manipulated (S, F, G and L plots), which strongly suggests that manipulating the order of arrival of forbs, grasses and legumes only had a weak effect on aboveground productivity in our experimental system (Figure 4; see Table S8 for statistics). Standing shoot biomass differed between free succession plots and other PFG order of arrival scenarios, but only in the second growing season of both sub-experiments. In that year, plots in which all functional groups were sown simultaneously were on average 118% (POEM2020) and 72% (POEM2021) more productive than free succession plots. In the first sub-experiment (POEM2020), grasses-first plots were also 129% more productive than free succession plots in the second growing season. In the same year, legumes-first plots were 69% more productive than free successions in the second sub-experiment (POEM2021). The year of initiation of each experiment also had a major impact on the aboveground productivity of plant communities. On average, plots were 38% less productive in the sub-experiment set up in 2021 than in the sub-experiment set up in 2020 (Figure 4, Table S8).

**Figure 4.**
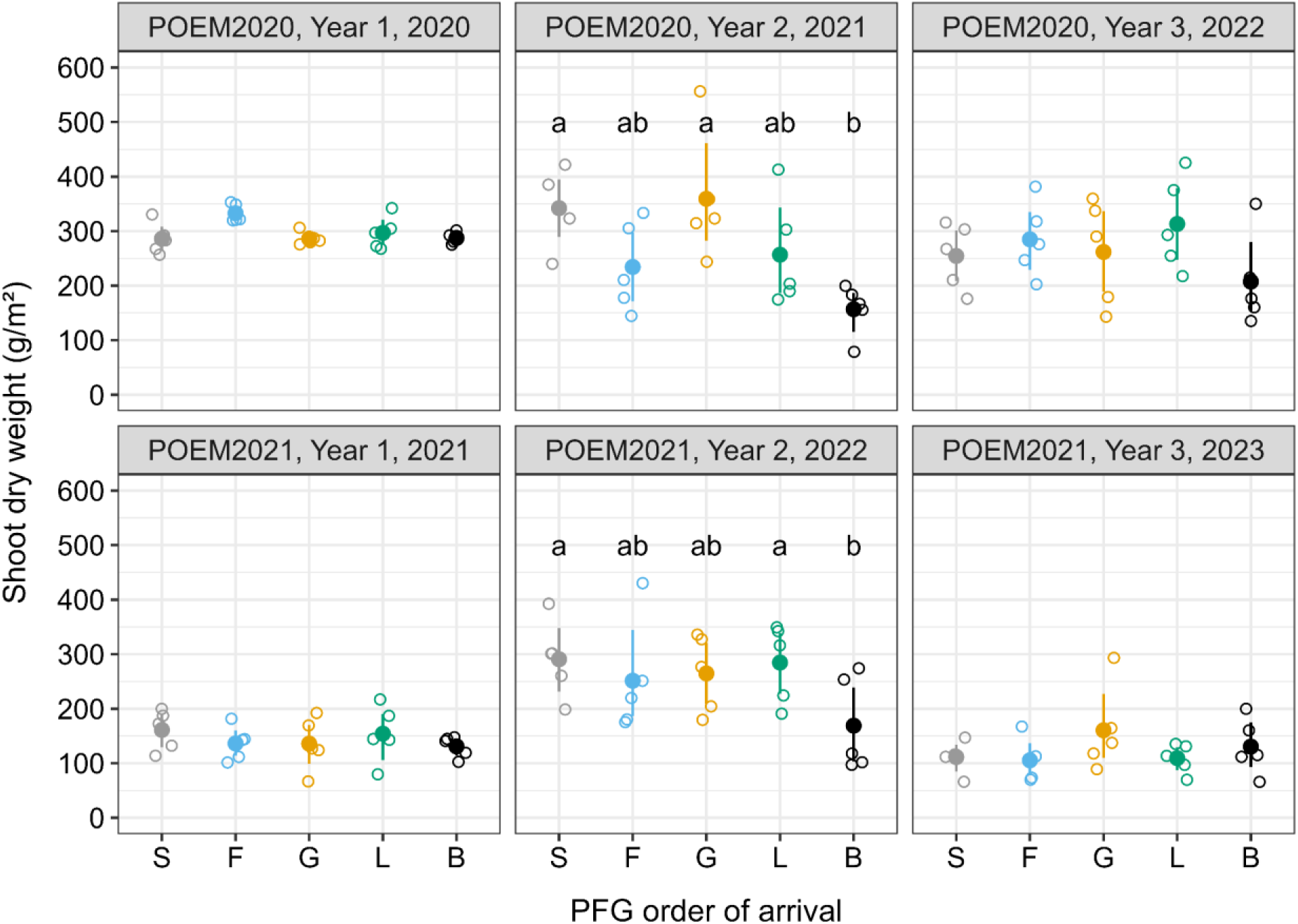
Year of initiation, not PFG order of arrival, is the main driver of aboveground productivity. For each combination of year of initiation, PFG of arrival and sampling year, the mean value (closed dot) and 95% confidence interval computed using non-parametric bootstrap are shown (n=5). Open dots represent the observed shoot dry weight values, which are jittered horizontally to improve readability. S, synchronous sowing of forbs, grasses and legumes; F, forbs sown first; G, grasses sown first; L, legumes sown first; B free succession plots.

### Root productivity was weakly affected by PFG order of arrival

Using the average planar root length density (*pRLD*, Figure 5) and average planar root surface density (*pRSD*, Figure S5) measured in a plot using minirhizotron images as proxies for standing root biomass production, we found that PFG order of arrival did not affect belowground productivity over the 3-year study period (*pRLD*: P = 0.275; *pRSD*: P = 0.166). At peak root production (day 772 on May 25, 2023), however, the planar root length density measured in synchronous plots was on average 13%, 24%, and 19% greater than in plots where forbs, grasses, or legumes were sown first.

**Figure 5.**
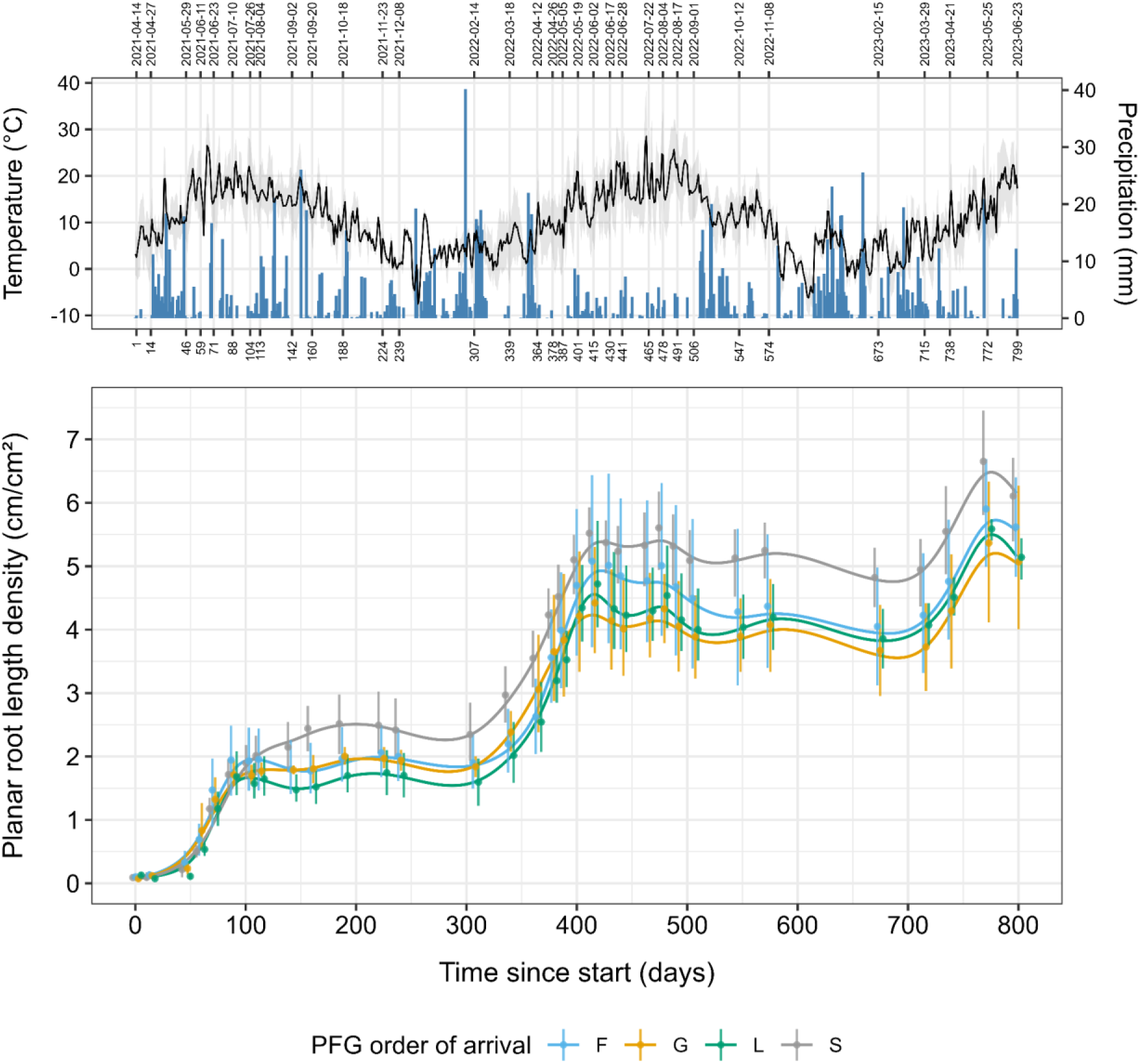
Root productivity was weakly affected by PFG order of arrival. Panel A shows the evolution of maximum, mean and minimum air temperature (°C), as well as daily precipitation (mm), at our experimental site between April 2021 and June 2023. Panel B shows the temporal evolution of the average planar root length density (*pRLD*) measured in POEM2021 plots for each PFG order of arrival scenario using minirhizotrons. Points and error bars indicate mean *pRLD* values and 95% confidence intervals (non-parametric bootstrap) measured at 33 time points spread over the first 800 days of POEM2021, respectively. Continuous lines are predictions from a generalised additive model. S, synchronous sowing of forbs, grasses and legumes; F, forbs sown first; G, grasses sown first; L, legumes sown first.

### Sowing forbs or legumes first led to deeper-rooted plant communities

Although our results do not provide strong support for the existence of differences in root productivity between PFG order of arrival scenarios, they do show that manipulating the arrival order of grasses, forbs and legumes had a strong impact on the vertical distribution of roots and the average rooting depth of plant communities (*pRLD*: P = 0.005; *pRSD*: P = 0.004). On average, forbs-first and legumes-first communities rooted 41% and 29% deeper than communities where grasses were sown first, respectively (Figure 6; F/G: P = 0.0002; G/L: P = 0.0109). We obtained similar results when the average rooting depth of plant communities was estimated using planar root surface density data (Figure S6; F/G: P = 0.0003; G/L: P = 0.0089). We did not find strong evidence to support that the average rooting depth in synchronous communities was different than in forbs-first, grasses-first or legumes-first communities (Figure 6; S/F: P = 0.076; S/G: P = 0.284; S/L: P = 0.542). Although all PFG order of arrival scenarios led to the accumulation of roots ~15 cm below the soil surface, we also observed an accumulation of roots deeper into the soil (~40-45 cm), but mostly in plots where forbs or legumes were sown first (Figures 6 and S6). This deeper root hotspot is particularly visible between 500 and 700 days after the start of POEM2021 (i.e., between September 2022 and March 2023).

**Figure 6.**
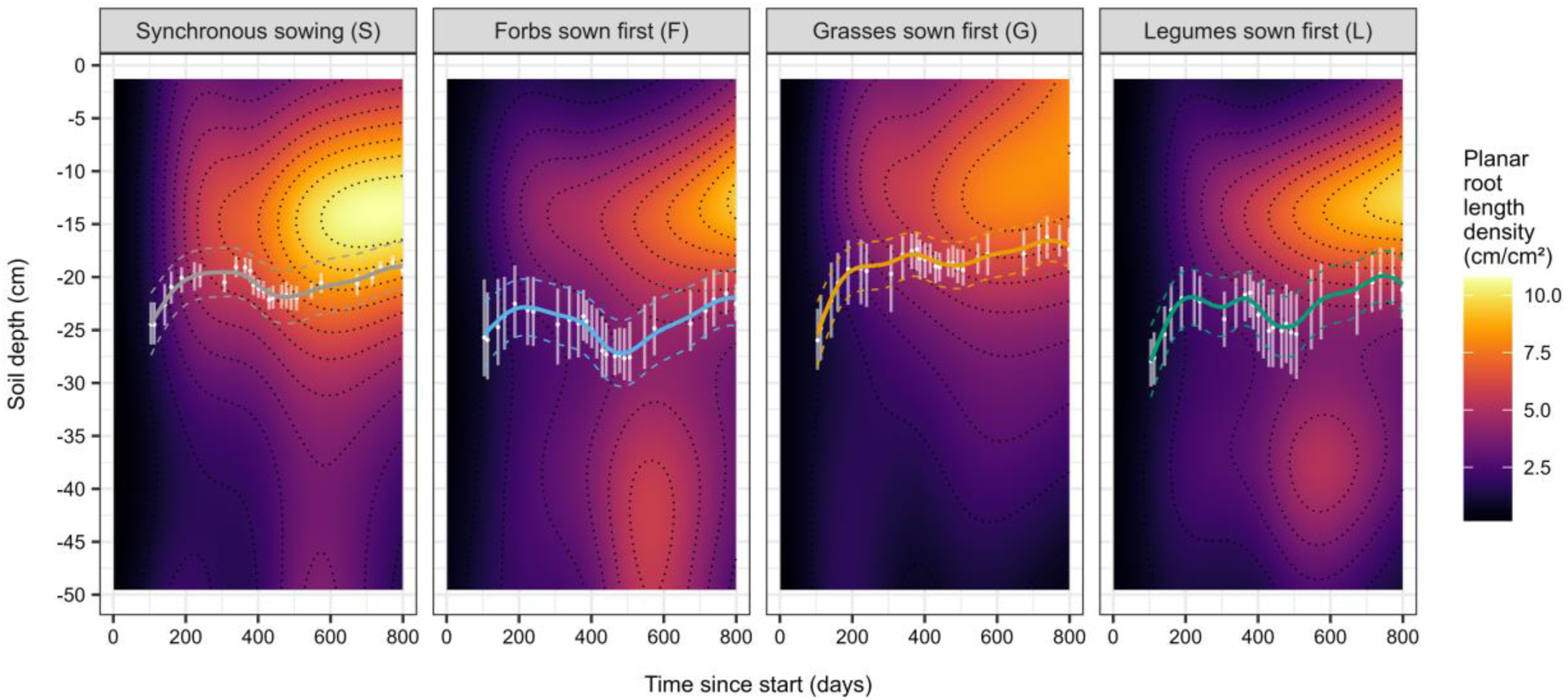
Sowing forbs or legumes first led to deeper-rooted plant communities. The raster image in each panel shows predictions from a first generalised additive model of the planar root length density (*pRLD*) as a function of time since the start of the experiment (0-800 days) and soil depth (1.4-49.5 cm). Results are plotted separately for each PFG order of arrival scenario. On top of each raster, white dots and error bars indicate the mean values and 95% confidence intervals (non-parametric bootstrap) of the mean rooting depth (*MRD*) estimated from planar root length density data between days 100 and 800, respectively. Continuous lines are predicted *MRD* values from a second generalised additive model.

## Discussion

The POEM experiment enabled us to quantify the relative importance of PFG order of arrival, year of initiation and time since establishment for the structure and functioning of grassland plant communities during the first three years of community assembly. Our results showed that the year of initiation had a strong impact on the aboveground productivity of plant communities, but was a less important driver of species composition and diversity than time since establishment and PFG order of arrival. Although PFG order of arrival did not affect aboveground and belowground productivity, our results demonstrate that it can modulate the vertical distribution of roots at the community level, with grasses-first communities rooting more shallowly than forbs-first and legumes-first communities.

Contrary to our first hypothesis, we found that species composition was mainly driven by time since establishment and, to a smaller extent, PFG order of arrival and year of initiation. Plant diversity, however, was dependent on PFG order of arrival, year of initiation, and time since establishment. We found that the effect of PFG order of arrival on plant species richness was dependent on time since establishment, but not year of initiation. Previous studies from mesotrophic grasslands have shown that priority effects can affect community composition, with groups of species arriving first dominating communities (Körner et al., 2008; von Gillhaussen et al., 2014; Weidlich et al., 2017), which we did not find as strongly in our study after three growing seasons. Our results are in line with the findings of prairie grassland experiments manipulating grass and forb order of arrival (Werner et al., 2016), except that order of arrival effects are weaker in our system, probably because of our shorter sowing interval between early- and late-arriving species (6 weeks vs 1 year) and the fact that we did not weed mixture plots after sowing. Year effects have also been documented as a factor affecting community composition (Groves & Brudvig, 2019; Werner et al., 2020). For example, in an experiment manipulating the timing of arrival of native and exotic grasses across three sites and four years, Stuble et al. (2017.b) found strong evidence that the strength of priority effects exerted by native species after one growing season when they were given a 2 week head start was modulated by site location and year of initiation. This contrasts with the results of our POEM experiment obtained at the end of the first growing season, which can certainly be explained by the fact that, unlike Stuble et al. (2017.b), we did not weed unsown species in the mixture plots in order to let plant communities undergo natural assembly processes. In our POEM experiment, the sown species required time to establish, during which unsown weedy agricultural species established themselves (from the seedbank) in the first growing season of both sub-experiments. These unsown species initially outcompeted the species sown in the plots. From the second growing season onwards, however, the sown species dominated the plots. This effect was probably influenced strongly by the timing of mowing in such dry acidic grasslands, which is later and less frequent (once versus twice per growing season) than in mesotrophic grasslands - as we know that mowing gives the perennial target species an advantage (Kirmer et al., 2018). Our findings are thus in line with expectations from ecological theory, that in more nutrient-poor dry acidic grasslands (e.g. our POEM experiment) competitive interactions (and hence asymmetric competition as a mechanism of priority effects within niche preemption) will be weaker (Chase, 2003), but also suggest that longer sowing intervals may be needed to create larger priority effects (Werner et al., 2016). In both POEM sub-experiments, the grasses-first treatment generally exhibited lower taxonomic diversity over the three growing seasons, possibly due to stronger competitive ability of grass species when given an initial advantage (Linder et al., 2018), allowing them to outcompete other species and create stronger priority effects (Cadotte, 2023; Werner et al., 2016).

Contrary to our second hypothesis, we did not find any strong effect of PFG order of arrival on standing shoot biomass production, with no difference in yield between synchronous, forbs-first, grasses-first, and legumes-first communities. However, we found that the year of initiation of an experiment was a stronger driver of aboveground community productivity, with plots sown in 2020 being on average more productive than those sown in 2021. The difference in productivity can probably be attributed to contrasting weather conditions during the first growing season of each sub-experiment (Atkinson et al., 2023; Bakker et al., 2003; Catano et al., 2023; Stuble et al., 2017.a; Werner et al., 2020), but more work is needed to identify key weather variables driving this year effect. In contrast to previous studies on priority effects that manipulated PFG order of arrival in grassland ecosystems (Körner et al., 2008; von Gillhaussen et al., 2014; Weidlich et al., 2017), our results did not support any difference in productivity between PFG order of arrival scenarios. The absence of PFG order of arrival effects on aboveground productivity in our study could be due to (1) unfavorable conditions for the establishment of the sown species, and (2) a too short sowing interval for nutrient-poor grasslands. For this reason, the next POEM sub-experiments will use a one year sowing interval between early- and late-arriving species groups.

Data from POEM2021, which is the only experiment so far equipped with minirhizotron tubes, did not support our third hypothesis. Using standing root length density (or root surface area) as a proxy, we did not find a strong effect of PFG order of arrival on root productivity (although synchronous plots tended to be more productive than the others). This contrasts with findings from previous studies manipulating PFG order of arrival that measured root productivity in containers (Körner et al., 2008) or in the topsoil under field conditions (Weidlich et al., 2018). These studies have shown that communities where legumes were sown first have a lower root productivity or lower standing root length density in the topsoil (Körner et al., 2008; Weidlich et al., 2018). This discrepancy may be due to methodological differences, e.g. in our study, root productivity was measured non-destructively over three years up to a depth of ~50 cm.

In agreement with our fourth hypothesis, we found that PFG order of arrival had a strong effect on vertical root distribution at the community level. Indeed, we found that sowing legumes or forbs first led to communities rooting deeper than when grasses were sown first. We found identical results when we manipulated PFG order of arrival in rhizoboxes under more controlled conditions (Alonso-Crespo et al., 2023). Considering that grassland species have different morphological and architectural characteristics, as well as different levels of root phenotypic plasticity in response to (a) biotic conditions (Bakker et al., 2019; Case et al., 2020; Chen et al., 2020; Herben et al., 2018; Lepik et al., 2021), slight differences in community composition following PFG order of arrival manipulation could lead to different patterns of root distribution. Given that grasses tend to root more superficially than forbs (Bakker et al., 2019, 2021; Chen et al., 2020), sowing grass species first may have increased root colonisation and interspecific competition in the topsoil, which could have made it more difficult for later-arriving forbs and legumes to grow in deeper soil layers, thus making root distribution of the entire community more shallow. The fact that data from this field experiment and another controlled experiment in rhizoboxes (Alonso-Crespo et al., 2023) support a strong effect of PFG order of arrival on root distribution without affecting root productivity strongly suggests that previous observations that legumes-first communities have a lower standing root length density in the topsoil (Weidlich et al., 2018) may actually be due to differences in vertical root distribution, and not to differences in root productivity. To better understand the mechanisms behind plant order of arrival effects on root distribution and their ecological consequences for plant communities, additional work is needed to measure root distribution at the species level, for instance using molecular techniques based on DNA sequencing (Wagemaker et al., 2020). In addition, a forthcoming second sub-experiment with minirhizotron measurements will allow us to see whether and how our root distribution findings are affected by year of initiation.

## Conclusions and outlook

In our study, we found that sowing legumes or forbs before the other plant functional groups caused deeper rooting than in other communities, whereas other factors such as year of initiation or time since establishment had a stronger effect on aboveground community structure (composition and diversity) and functioning (aboveground biomass production). This is one of the few studies that experimentally looked at priority and year effects on both aboveground and belowground dynamics and our findings have potentially important implications for grassland restoration. If our findings can be generalised for a range of different grassland types, then sowing legumes or forbs-first could create communities with deeper roots that are more species-rich. This in turn could lead to plant communities that are more resistant/resilient to extreme weather events (e.g. drought) and potentially could improve soil carbon storage at depth. In order for our findings to be useful for grassland restoration, we need to better understand the mechanisms that determine the effects of plant order of arrival on root distribution, as well as identify the key environmental variables that drive the year effects that we found.

## Acknowledgments

This work was funded by the German Research Foundation (project number: 420444099). The authors would like to thank Christoph Stegen for his exceptional technical support. Many thanks to all the colleagues and student helpers who helped us set up and maintain the POEM experiment.

## Author Contributions

VMT, BMD and MS conceived the project, designed the experiment and secured funding; IMAC, BMD, TN and VMT collected data; IMAC and BMD analysed root images; IMAC, BMD and AF analysed data; IMAC produced the first draft of the manuscript, with support from BMD, AF and VMT. All authors contributed critically to the drafts and gave final approval for publication.

## Data availability statement

The data and R codes that support the findings of this study are openly available in Zenodo at https://doi.org/10.5281/zenodo.10119982.

