## Supplementary information for "Exploring priority and year effects on plant diversity, productivity and vertical root distribution: first insights from a grassland field experiment"

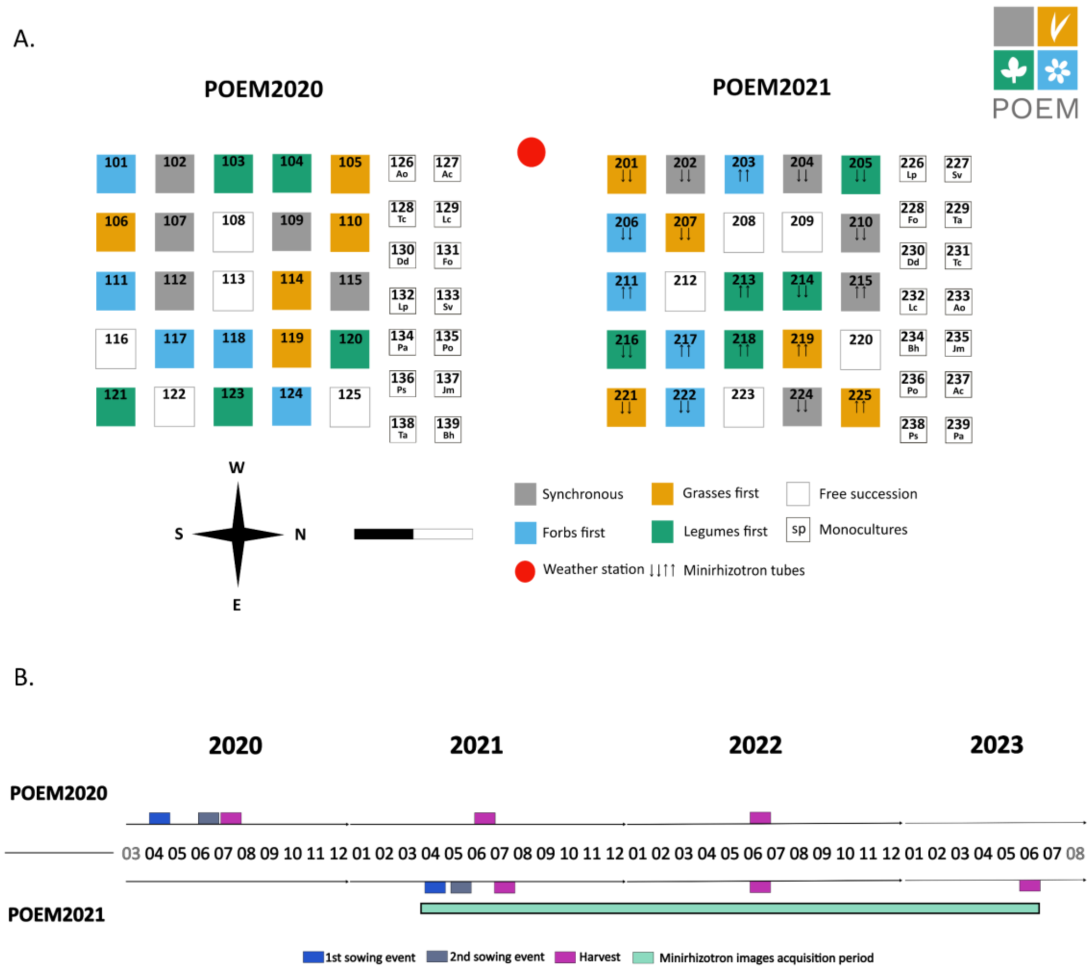


**Figure S1. Overview of the POEM experiment. A. Experimental design.** As of June 2023, the experiment consists of two sub-experiments set up in different years: POEM2020 (set up in 2020) and POEM2021 (set up in 2021). Each sub-experiment consists of 25 mixture plots (3×3 m²) and 14 monoculture plots (3×3 m²). Each PFG order of arrival scenario is represented by 5 replicates. Ac, *Agrostis capillaris*; Ao, *Anthoxanthum odoratum*; Bh, *Bromus hordeaceus*; Dd, *Dianthus deltoides*; Fo, *Festuca ovina*; Jm, *Jasione montana*; Lp, *Lathyrus pratensis*; Lc, *Lotus corniculatus*; Po, *Pilosella officinarum*; Ps, *Pimpinella saxifraga*; Pa, *Potentilla argentea*; Sv, *Silene vulgaris*; Ta, *Trifolium arvense*, Tc, *Trifolium campestre*. **B. Experimental timeline**. This timeline includes activities carried out in POEM2020 and POEM2021 during the first three growing seasons.

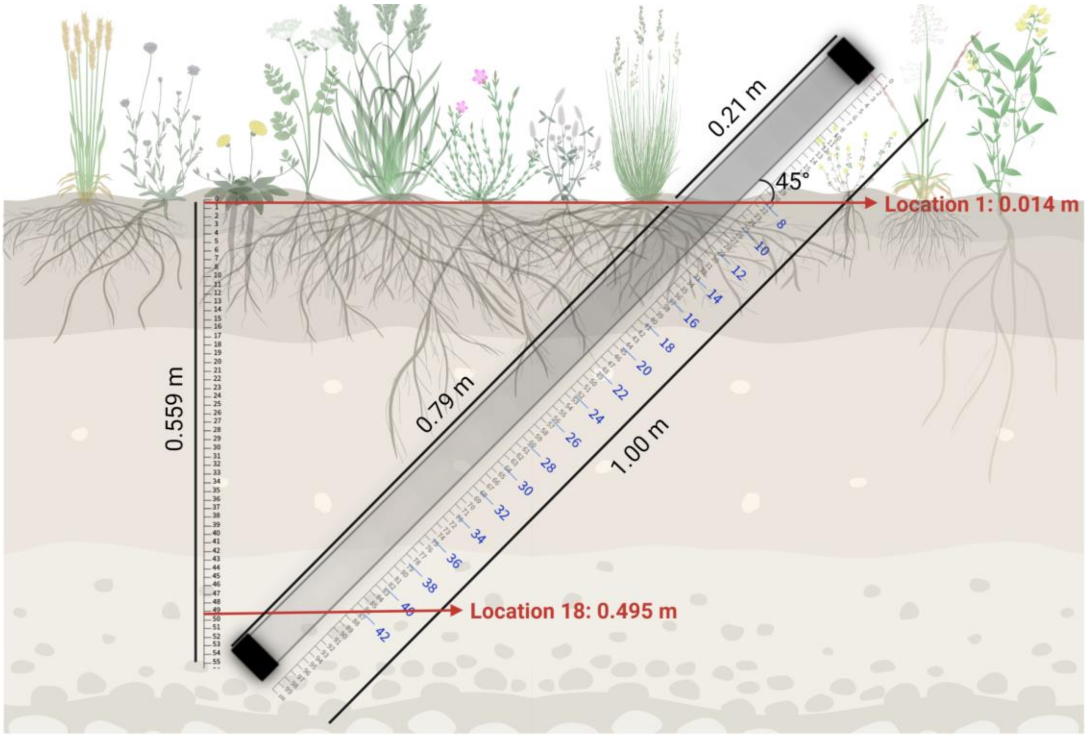


**Figure S2.** Installation of minirhizotron tubes in POEM2021 plots. Minirhizotron tubes were installed at a 45° angle. A step-by-step description of the installation of minirhizotron tubes in POEM2021 is available in the video provided as supplementary material. Roots growing along a minirhizotron tube were regularly imaged at 18 equally spaced locations (in blue) along the tube. Using the setup shown in the figure, root images were collected between 1.4 cm depth and 49.5 cm depth.

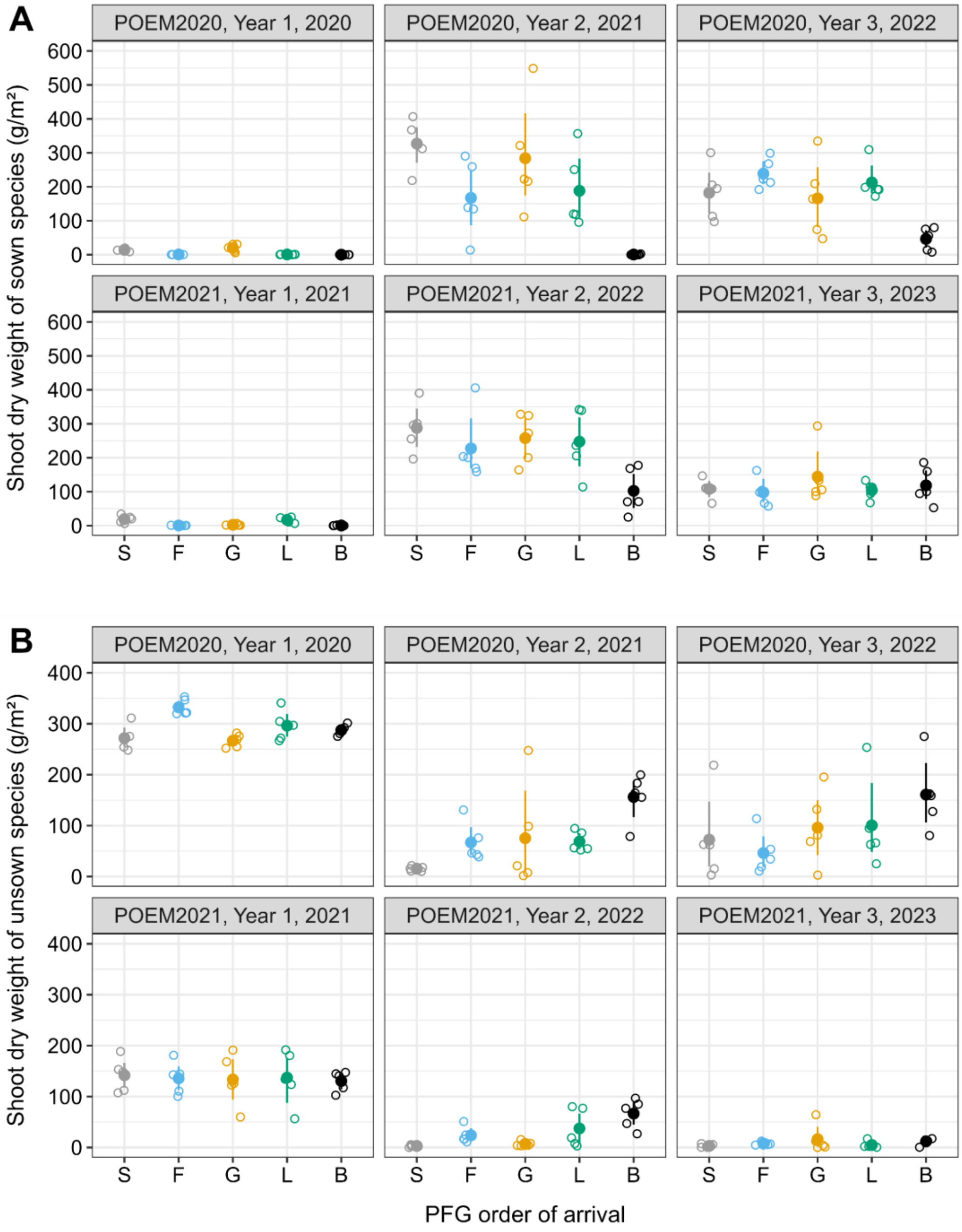


**Figure S3. Effects of year of initiation, sampling year and PFG order of arrival on the aboveground productivity of sown (A) and unsown (B) species.** For each combination of year of initiation, PFG of arrival and sampling year, the mean value (closed dot) and 95% confidence interval computed using non-parametric bootstrap are shown (*n*=5). Open dots represent the observed shoot dry weight values, which are jittered horizontally to improve readability. S, synchronous sowing of forbs, grasses and legumes; F, forbs sown first; G, grasses sown first; L, legumes sown first; B free succession plots.

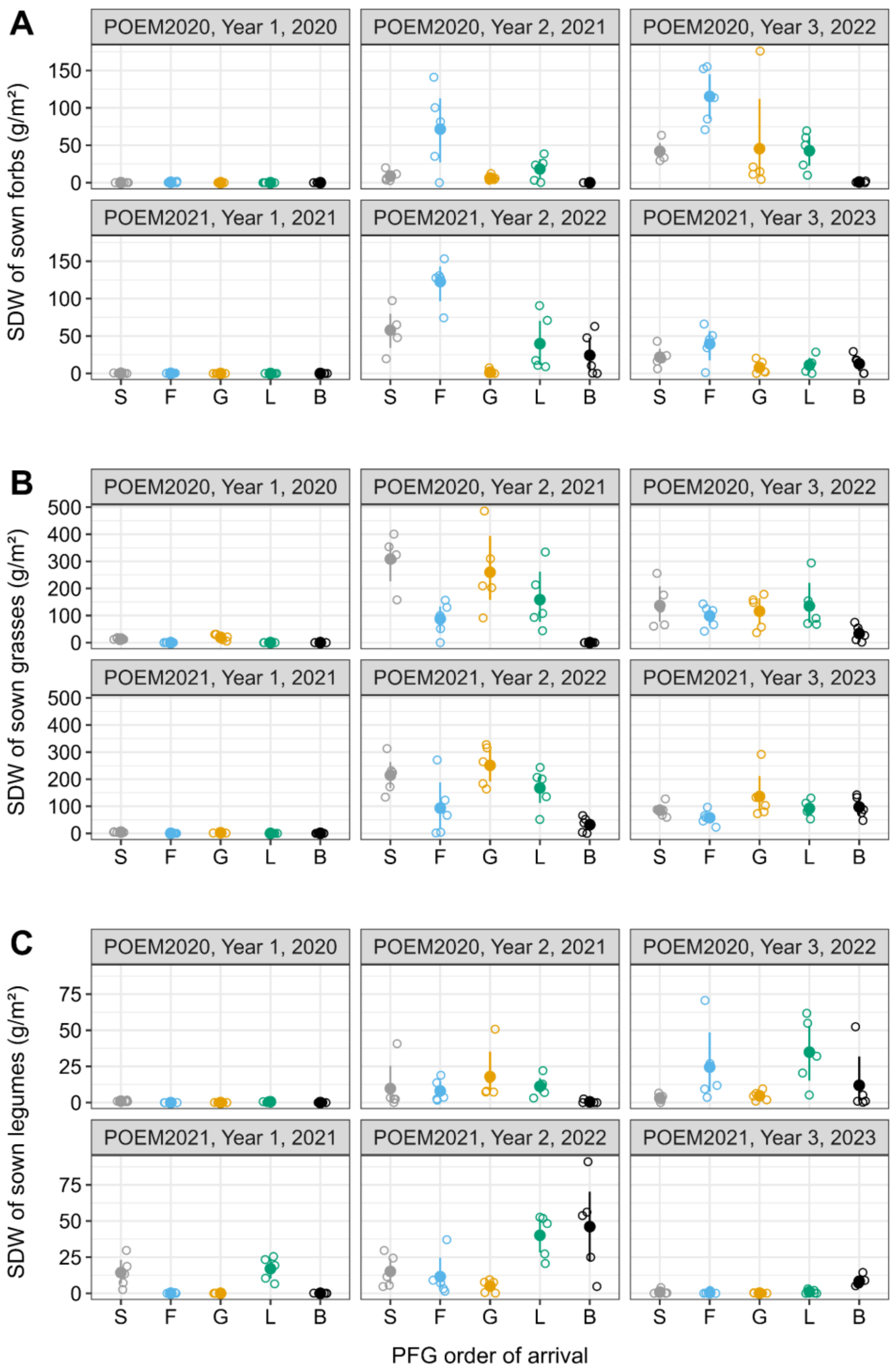


**Figure S4. Effects of year of initiation, sampling year and PFG order of arrival on the aboveground productivity of sown forbs (A), grasses (B) and legumes (C).** For each combination of year of initiation, PFG of arrival and sampling year, the mean value (closed dot) and 95% confidence interval computed using non-parametric bootstrap are shown (n=5). Open dots represent the observed shoot dry weight values, which are jittered horizontally to improve readability. S, synchronous sowing of forbs, grasses and legumes; F, forbs sown first; G, grasses sown first; L, legumes sown first; B free succession plots.

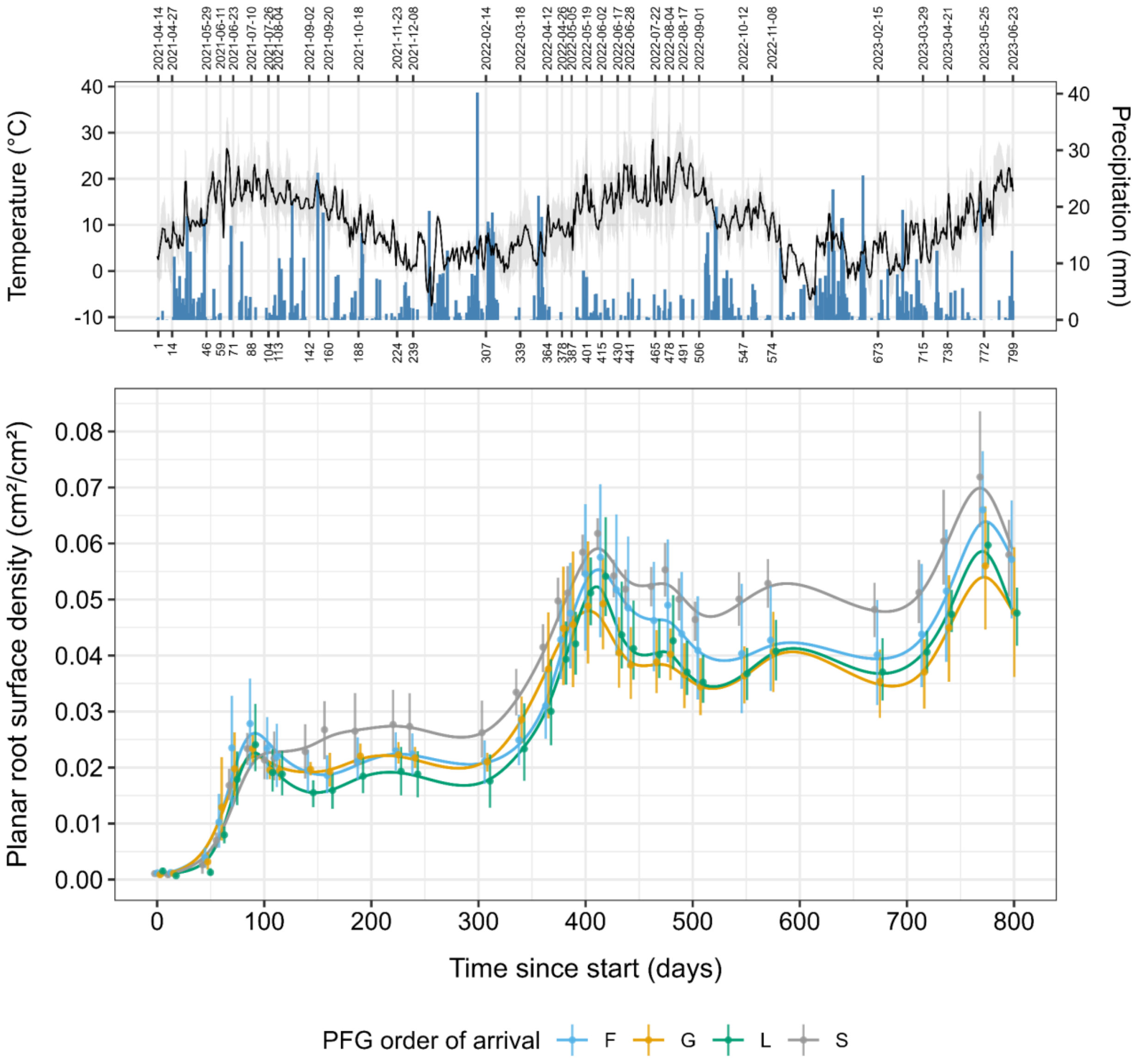


**Figure S5. Root productivity was weakly affected by PFG order of arrival. Panel A shows the evolution of maximum, mean and minimum air temperature (°C), as well as daily precipitation (mm), at our experimental site between April 2021 and June 2023.** Panel B shows the temporal evolution of the average planar root surface density (*pRSD*) measured in POEM2021 plots for each PFG order of arrival scenario using minirhizotrons. Points and error bars indicate mean *pRSD* values and 95% confidence intervals (non-parametric bootstrap) measured at 33 time points spread over the first 800 days of POEM2021, respectively. Continuous lines are predictions from a generalised additive model. S, synchronous sowing of forbs, grasses and legumes; F, forbs sown first; G, grasses sown first; L, legumes sown first.

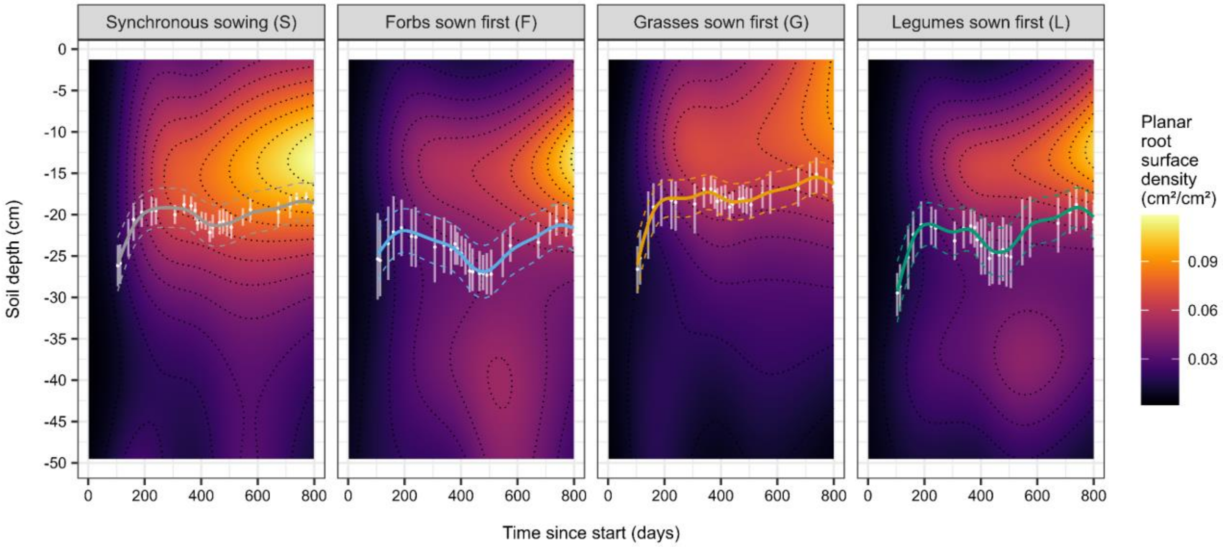


**Figure S6. Sowing forbs or legumes first led to deeper-rooted plant communities.** The raster image in each panel shows predictions from a first generalised additive model of the planar root surface density (*pRSD*) as a function of time since the start of the experiment (0-800 days) and soil depth (1.4-49.5 cm). Results are plotted separately for each PFG order of arrival scenario. On top of each raster, white dots and error bars indicate the mean values and 95% confidence intervals (non-parametric bootstrap) of the mean rooting depth (*MRD*) estimated from planar root surface density data between days 100 and 800, respectively. Continuous lines are predicted *MRD* values from a second generalised additive model.

**Table S1. Composition of the seed mixtures used in the mixture plots of POEM2020 and POEM2021**. For each species, the target number of individuals in a plot was calculated considering (1) the plot surface area (9 m²), (2) the desired plant density (1000 plants/m²), (3) the fact that PFGs should be equally represented in the seed mixture, and (4) the fact that species within each PFG should be equally represented in the seed mixture. 'Seeds per plot' is the total number of seeds needed to reach the target number of individuals of a species in a plot. 'Seed mass per plot' is the corresponding seed mass value. TSW, thousand seed weight (g); FG, functional group (F for forbs, G for grasses, L for legumes); GR, germination rate [0,1].

| **Species** | **FG** | **Target number of ind per plot** | **POEM2020** | | | | **POEM2021** | | | |
|---|---|---|---|---|---|---|---|---|---|---|
| | | | TSW (g) | GR | Seeds per plot | Seed mass per plot (g) | TSW (g) | GR | Seeds per plot | Seed mass per plot (g) |
| *Lotus corniculatus* | L | 750 | 1.25 | 0.71 | 1050 | 1.31 | 1.36 | 0.76 | 987 | 1.34 |
| *Trifolium arvense* | L | 750 | 0.31 | 0.31 | 2386 | 0.74 | 0.43 | 0.51 | 1463 | 0.63 |
| *Trifolium campestre* | L | 750 | 0.49 | 0.31 | 2386 | 1.18 | 0.53 | 0.56 | 1339 | 0.71 |
| *Lathyrus pratensis* | L | 750 | 10.65 | 0.51 | 1458 | 15.54 | 9.40 | 0.68 | 1099 | 10.33 |
| *Anthoxanthum odoratum* | G | 750 | 0.51 | 0.40 | 1875 | 0.95 | 0.47 | 0.48 | 1579 | 0.74 |
| *Bromus hordeaceus* | G | 750 | 2.55 | 0.74 | 1010 | 2.57 | 2.70 | 0.52 | 1442 | 3.89 |
| *Festuca ovina* | G | 750 | 0.28 | 0.31 | 2386 | 0.67 | 0.25 | 0.16 | 4688 | 1.18 |
| *Agrostis capillaris* | G | 750 | 0.07 | 0.86 | 875 | 0.06 | 0.09 | 0.28 | 2727 | 0.25 |
| *Silene vulgaris* | F | 500 | 0.59 | 0.77 | 648 | 0.38 | 0.55 | 0.88 | 568 | 0.31 |
| *Dianthus deltoides* | F | 500 | 0.16 | 0.89 | 565 | 0.09 | 0.18 | 0.92 | 543 | 0.10 |
| *Pilosella officinarum* | F | 500 | 0.19 | 0.76 | 658 | 0.12 | 0.19 | 0.92 | 543 | 0.10 |
| *Jasione montana* | F | 500 | 0.02 | 0.74 | 673 | 0.01 | 0.02 | 0.50 | 1000 | 0.02 |
| *Potentilla argentea* | F | 500 | 0.13 | 0.51 | 972 | 0.12 | 0.07 | 0.24 | 2083 | 0.16 |
| *Pimpinella saxifraga* | F | 500 | 0.73 | 0.13 | 3846 | 2.81 | 0.93 | 0.20 | 2500 | 2.32 |

**Table S2. Seed mass used in the monoculture plots of POEM2020 and POEM2021.** The target number of individuals in a plot was calculated considering (1) the plot surface area (4 $m^2$) and (2) the desired plant density (1000 plants/$m^2$). 'Seeds per plot' is the total number of seeds needed to reach the target number of individuals of a species in a plot. 'Seed mass per plot' is the corresponding seed mass value. TSW, thousand seed weight (g); FG, functional group (F for forbs, G for grasses, L for legumes); GR, germination rate [0,1].

| **Species** | **FG** | **Target number of ind per plot** | **POEM2020** | | | | **POEM2021** | | | |
|---|---|---|---|---|---|---|---|---|---|---|
| | | | TSW (g) | GR | Seeds per plot | Seed mass per plot (g) | TSW (g) | GR | Seeds per plot | Seed mass per plot (g) |
| *Lotus corniculatus* | L | 4000 | 1.25 | 0.71 | 5600 | 6.97 | 1.36 | 0.76 | 5263 | 7.16 |
| *Trifolium arvense* | L | 4000 | 0.31 | 0.31 | 12727 | 3.95 | 0.43 | 0.51 | 7805 | 3.35 |
| *Trifolium campestre* | L | 4000 | 0.49 | 0.31 | 12727 | 6.27 | 0.53 | 0.56 | 7143 | 3.78 |
| *Lathyrus pratensis* | L | 4000 | 10.65 | 0.51 | 7778 | 82.87 | 9.40 | 0.68 | 5861 | 55.09 |
| *Anthoxanthum odoratum* | G | 4000 | 0.51 | 0.40 | 10000 | 5.08 | 0.47 | 0.48 | 8421 | 3.97 |
| *Bromus hordeaceus* | G | 4000 | 2.55 | 0.74 | 5385 | 13.72 | 2.70 | 0.52 | 7692 | 20.75 |
| *Festuca ovina* | G | 4000 | 0.28 | 0.31 | 12727 | 3.58 | 0.25 | 0.16 | 25000 | 6.29 |
| *Agrostis capillaris* | G | 4000 | 0.07 | 0.86 | 4667 | 0.33 | 0.09 | 0.28 | 14545 | 1.34 |
| *Silene vulgaris* | F | 4000 | 0.59 | 0.77 | 5185 | 3.07 | 0.55 | 0.88 | 4545 | 2.48 |
| *Dianthus deltoides* | F | 4000 | 0.16 | 0.89 | 4516 | 0.72 | 0.18 | 0.92 | 4348 | 0.77 |
| *Pilosella officinarum* | F | 4000 | 0.19 | 0.76 | 5263 | 0.98 | 0.19 | 0.92 | 4348 | 0.81 |
| *Jasione montana* | F | 4000 | 0.02 | 0.74 | 5385 | 0.10 | 0.02 | 0.50 | 8000 | 0.18 |
| *Potentilla argentea* | F | 4000 | 0.13 | 0.51 | 7778 | 0.97 | 0.07 | 0.24 | 16667 | 1.24 |
| *Pimpinella saxifraga* | F | 4000 | 0.73 | 0.13 | 30769 | 22.46 | 0.93 | 0.20 | 20000 | 18.60 |

**Table S3. Results of permutational multivariate analysis of variance (PERMANOVA) testing the effects of PFG order of arrival, year of initiation and sampling year on dissimilarities in plant species composition using species-specific biomass data.** The Bray-Curtis dissimilarity index was used. Terms were added sequentially (first to last). Number of permutations: 1000. See also figure 2A.

| | Df | Sum of squares | $R^2$ | F | P-value |
|---|---|---|---|---|---|
| **PFG order of arrival (a)** | 4 | 3.46 | **0.077** | 7.94 | **0.000999** |
| **Sampling year (b)** | 2 | 17.70 | **0.393** | 81.22 | **0.000999** |
| **Year of initiation (c)** | 1 | 2.61 | **0.058** | 23.97 | **0.000999** |
| **a*b** | 8 | 2.79 | **0.062** | 3.20 | **0.000999** |
| **a*c** | 4 | 1.09 | **0.024** | 2.50 | **0.002997** |
| **b*c** | 2 | 3.11 | **0.069** | 14.27 | **0.000999** |
| a*b*c | 8 | 1.15 | 0.026 | 1.32 | 0.081918 |
| Residual | 120 | 13.07 | 0.291 | | |
| Total | 149 | 44.98 | 1.000 | | |

**Table S4. Results of permutational multivariate analysis of variance (PERMANOVA) testing the effects of PFG order of arrival, year of initiation and sampling year on dissimilarities in plant species composition using presence/absence data.** The Jaccard dissimilarity index was used. Terms were added sequentially (first to last). Number of permutations: 1000. See also figure 2B.

| | Df | Sum of squares | $R^2$ | F | P-value |
|---|---|---|---|---|---|
| **PFG order of arrival (a)** | 4 | 1.75 | **0.097** | 8.23 | **0.000999** |
| **Sampling year (b)** | 2 | 6.76 | **0.373** | 63.50 | **0.000999** |
| **Year of initiation (c)** | 1 | 1.48 | **0.081** | 27.72 | **0.000999** |
| **a*b** | 8 | 1.03 | **0.057** | 2.42 | **0.000999** |
| **a*c** | 4 | 0.52 | **0.029** | 2.47 | **0.000999** |
| **b*c** | 1 | 1.12 | **0.062** | 21.03 | **0.000999** |
| a*b*c | 4 | 0.12 | 0.007 | 0.59 | 0.966034 |
| Residual | 100 | 5.32 | 0.294 | | |
| Total | 124 | 18.11 | 1.000 | | |

**Table S5. Results of a generalised linear mixed-effect model testing the effects of PFG order of arrival, year of initiation and sampling year on the effective number of species at *q*=0 (species richness).** Plot ID was used as a random effect in the model (variance: 0.0049). The model was fitted using a Gamma distribution and a log-link function. This analysis of deviance table (Type III Wald chi square tests) was produced using the Anova function in the car package (Fox and Weisberg, 2019). See also Figure 3.

| | Chisq | Df | P-value |
|---|---|---|---|
| (Intercept) | 1027.20 | 1 | **< 2.2e-16** |
| **PFG order of arrival (a)** | 22.53 | 4 | **0.0002** |
| **Sampling year (b)** | 6.91 | 2 | **0.0316** |
| Year of initiation (c) | 0.96 | 1 | 0.3262 |
| **a*b** | 24.52 | 8 | **0.0019** |
| a*c | 5.03 | 4 | 0.2843 |
| b*c | 3.41 | 2 | 0.1817 |
| a*b*c | 6.01 | 8 | 0.6461 |

**Table S6. Results of a generalised linear mixed-effect model testing the effects of PFG order of arrival, year of initiation and sampling year on the effective number of species at *q*=1.** Plot ID was used as a random effect in the model (variance: 0.0127). The model was fitted using a Gamma distribution and a log-link function. This analysis of deviance table (Type III Wald chi square tests) was produced using the Anova function in the car package (Fox and Weisberg, 2019). See also Figure 3.

| | Chisq | Df | P-value |
|---|---|---|---|
| (Intercept) | 111.93 | 1 | **< 2.2e-16** |
| PFG order of arrival (a) | 3.93 | 4 | 0.4149 |
| **Sampling year (b)** | 27.15 | 2 | **1.27e-06** |
| Year of initiation (c) | 0.11 | 1 | 0.7370 |
| **a*b** | 73.79 | 8 | **8.62e-13** |
| a*c | 6.59 | 4 | 0.1591 |
| **b*c** | 14.21 | 2 | **0.0008** |
| **a*b*c** | 16.97 | 8 | **0.0304** |

**Table S7. Results of a generalised linear mixed-effect model testing the effects of PFG order of arrival, year of initiation and sampling year on the effective number of species at *q*=2.** Plot ID was used as a random effect in the model (variance: 0.0174). The model was fitted using a Gamma distribution and a log-link function. This analysis of deviance table (Type III Wald chi square tests) was produced using the Anova function in the car package (Fox and Weisberg, 2019). See also Figure 3.

| | Chisq | Df | P-value |
|---|---|---|---|
| (Intercept) | 56.18 | 1 | **6.60e-14** |
| PFG order of arrival (a) | 4.06 | 4 | 0.3979 |
| **Sampling year (b)** | 23.68 | 2 | **7.21e-06** |
| Year of initiation (c) | 0.83 | 1 | 0.3628 |
| **a*b** | 61.76 | 8 | **2.10e-10** |
| a*c | 4.81 | 4 | 0.3073 |
| **b*c** | 15.74 | 2 | **0.0004** |
| a*b*c | 15.33 | 8 | 0.0530 |

**Table S8. Results of a generalised linear mixed-effect model testing the effects of PFG order of arrival, year of initiation and sampling year on total shoot dry weight ($g/m^2$).** Plot ID was used as a random effect in the model (variance: 0.0091). The model was fitted using a Gamma distribution and a log-link function. This analysis of deviance table (Type III Wald chi square tests) was produced using the Anova function in the car package (Fox and Weisberg, 2019). See also Figure 4.

| | Chisq | Df | P-value |
|---|---|---|---|
| (Intercept) | 1963.29 | 1 | **< 2.2e-16** |
| PFG order of arrival (a) | 1.01 | 4 | 0.9077 |
| Sampling year (b) | 3.58 | 2 | 0.1669 |
| **Year of initiation (c)** | 9.97 | 1 | **0.0016** |
| **a*b** | 24.85 | 8 | **0.0016** |
| a*c | 1.96 | 4 | 0.7433 |
| **b*c** | 9.04 | 2 | **0.0109** |
| **a*b*c** | 16.90 | 8 | **0.0312** |
